# Nuclear decoupling is part of a rapid protein-level cellular response to high-intensity mechanical loading

**DOI:** 10.1101/317404

**Authors:** Hamish T.J. Gilbert, Venkatesh Mallikarjun, Oana Dobre, Mark R. Jackson, Robert Pedley, Andrew P. Gilmore, Stephen M. Richardson, Joe Swift

**Affiliations:** Wellcome Trust Centre for Cell-Matrix Research.; Division of Cell Matrix Biology and Regenerative Medicine, School of Biological Sciences.; Division of Molecular and Clinical Cancer Sciences, School of Medical Sciences.; Faculty of Biology, Medicine and Health, Manchester Academic Health Science Centre, University of Manchester, Manchester, M13 9PL, UK.; Present address: School of Engineering, University of Glasgow, Glasgow, G12 8QQ, UK.; Present address: Institute of Cancer Sciences, Glasgow, G61 1QH, UK.; Correspondence should be addressed to J.S.

## Abstract

Our current understanding of cellular mechano-signalling is based on static models, which do not replicate the dynamics of living tissues. Here, we examined the time-dependent response of primary human mesenchymal stem cells (hMSCs) to cyclic tensile strain (CTS). At low-intensity strain (1 hour, 4% CTS at 1 Hz) morphological changes mimicked responses to increased substrate stiffness. As the strain regime was intensified (frequency increased to 5 Hz), we characterised rapid establishment of a broad, structured and reversible protein-level response, even as transcription was apparently downregulated. Protein abundance was quantified coincident with changes to protein conformation and post transcriptional modification. Furthermore, we characterised changes within the linker of nucleo- and cytoskeleton (LINC) complex of proteins that bridges the nuclear envelope, and specifically to levels of SUN domain-containing protein 2 (SUN2). The result of this regulation was to decouple mechano-transmission between the cytoskeleton and the nucleus, thus conferring protection to chromatin.

---

The stiffness of tissue correlates with its ability to resist mechanical damage, with the structure and integrity of the human body defined by stiff tissues such as skin, muscle, cartilage and bone. Tissue mechanical properties are determined by the extracellular matrix (ECM), in particular by the identities and concentrations of its constitutive proteins^1-3^. ECM properties are further modulated by protein cross-linking, post-translational modifications (PTMs) and higher-order organisation. Cells resident within tissues maintain mechanical equilibrium with their environments^4, 5^, and the mechanical properties of cells are also regulated by the identities, concentrations, conformations and PTMs of structural intracellular proteins^1, 3, 6, 7^. The characteristics of adherent cells can be influenced by physical stimulation from the surrounding ECM, with protein content^1^, morphology^1, 8^, motility^9, 10^ and differentiation potential^11, 12^ amongst behaviours known to be affected by stiffness. Cells in living tissues experience microenvironments of diverse stiffness^5^, but are also subject to deformation during activity. Cells sense and respond to mechanical signals through pathways of mechanotransduction^13-15^, but must also maintain integrity and homeostasis within the tissue. A mismatch between mechanical loading and cellular regulation can contribute to pathology, such as in musculoskeletal and connective tissue disorders^16^, with ageing being a significant risk factor^17^.

Here, we sought to compare responses to stiffness and mechanical loading in primary human mesenchymal stem cells (hMSCs), a cell type with important physiological and reparative roles, that have led to investigations of their therapeutic potential in tissues such as muscle^18^ and heart^19^. We contrast cellular responses to stiffness and strain cycling, and identify a rapid, reversible and structured regulation of the proteome following high-intensity mechanical loading. Furthermore, we identify SUN domain-containing protein 2 (SUN2) as a strain-induced breakpoint in the linker of nucleo- and cytoskeleton (LINC) complex of proteins that acts as a pathway of intracellular mechano-transmission^13, 20^, thus enabling the nucleus to ‘decouple’ from the cytoskeleton in response to intense strain.

## RESULTS

### Cyclic tensile strain (CTS) uncouples the correlation between cellular and nuclear morphology

Primary hMSCs were cultured on stiffness-controlled polyacrylamide hydrogels or silicone elastomer sheets that could be subjected to CTS (both collagen-I coated). hMSCs were found to spread increasingly on stiffer substrates over a physiological range (2 – 50 kPa, cultured for 3 days; Fig. 1a, Supplementary Fig. 1a), as has been reported previously^1, 21^. Cells subjected to sinusoidal, equiaxial CTS for 1 hour at 1 or 2 Hz (change in strain = 4%) showed significantly increased spreading immediately after loading (*p* ≤ 0.05), returning to initial spread areas after 24 hours (Fig. 1b, Supplementary Figs. 1b, c). Earlier reports of cell behaviour following strain have described cell alignment relative to the direction of strain^22, 23^ and reorganization of focal adhesion (FA) complexes and the cytoskeleton^24-26^. As the strain applied in our system had radial symmetry, no overall alignment was observed, but increased cell spreading was consistent with previous reports describing FA activation^27^. The increase in spreading of hMSCs following dynamic straining at 1 and 2 Hz was thus similar to that observed with changes in static substrate stiffness.

**Figure 1.**
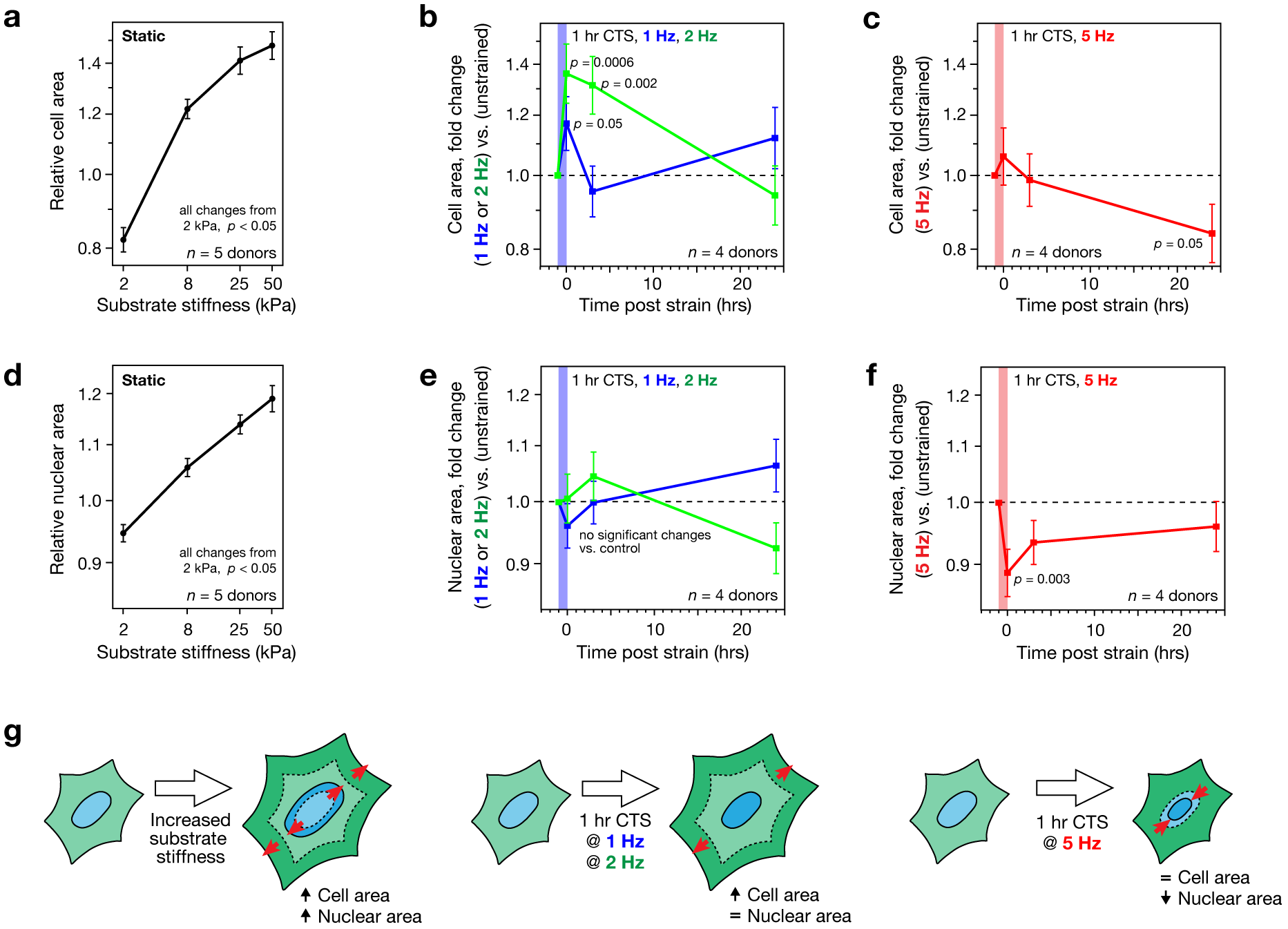
Loss of coupled cell and nuclear morphologies in primary human mesenchymal stem cells (hMSCs) following cyclic tensile strain (CTS). (**a**) Relative spreading areas of hMSCs cultured on static substrates (collagen-I coated polyacrylamide hydrogels; 2 – 50 kPa; *n* = 5 donors). Cell areas were found to increase as a function of substrate stiffness. (**b**) Cell areas of hMSCs following low-intensity CTS (0 – 4% strain at 1 or 2 Hz for 1 hour; *n* = 4 donors), normalized to unstrained controls. Cell areas increased immediately following strain, returning to pre-strained values after 3 hours in the 1 Hz treated cells, and after 24 hrs in the 2 Hz treated cells. (**c**) Cell areas of hMSCs following high-intensity CTS (2.6 – 6.2% strain at 5 Hz for 1 hour; *n* = 4 donors), normalised to unstrained controls. Cell areas remained unchanged immediately and 3 hours following strain treatment, but decreased 24 hours post-strain. Cell area normalisation was matched across figure parts (a), (b) and (c). (**d**) Relative nuclear areas of hMSCs cultured on static substrates (collagen-I coated PA hydrogels; 2 – 50 kPa; *n* = 5 donors). Nuclear areas increased with substrate stiffness, mirroring increased cell spreading. (**e**) Nuclear areas of hMSCs after low-intensity CTS (0 – 4% strain at 1 or 2 Hz for 1 hour; *n* = 4 donors), normalized to unstrained controls. Nuclear areas remained unchanged following strain treatment at 1 or 2 Hz. (**f**) Nuclear areas of hMSCs cultured following high-intensity CTS (2.6 – 6.2% strain at 5 Hz for 1 hour; *n* = 4 donors), normalised to unstrained controls. Nuclear areas decreased immediately following strain treatment, returning to pre-strained values after 3 hours. Nuclear area normalisation was matched across figure parts (d), (e) and (f). Data displayed as mean ± s.e.m.; *p*-values were determined from linear modeling. See Supplementary Figs. 1a, b for example cell images and Supplementary Figs. 1c, f for an examination of donor-to-donor variability. (**g**) Cartoon summarizing changes to cell and nuclear morphologies on stiffness-controlled substrates and immediately following 1 hour CTS. Cell and nuclear areas appeared coupled on increasingly stiff substrates. Low intensity CTS resulted in increased cell area, but no change in nuclear area; high-intensity CTS caused nuclear area to decrease independently of cell area.

To explore mechanisms that allow cells to endure more challenging mechanical environments, we increased the frequency of CTS to 5 Hz (change in strain = 3.6%; referred to henceforth as ‘high-intensity CTS’). The increased cell spreading observed at lower frequencies was not seen following 1 hour of high-intensity CTS (Fig. 1c). Cell spreading was significantly decreased 24 hours after treatment (*p* = 0.05), but cells remained attached to the substrate. Furthermore, neither cell viability nor proliferation were significantly affected (Supplementary Figs. 1d, e).

The nuclear area of hMSCs was found to increase with cell spreading on stiffer substrates (Fig. 1d, Supplementary Fig. 1a). This agrees with findings in earlier works^1, 28^ and reflects the interconnected nature of the cyto- and nucleoskeleton^29^, which has been shown to be necessary for mechanotransduction^20^. However, we found that this correlated behaviour of cell and nuclear spreading was lost in hMSCs subjected to CTS: there were no significant changes in nuclear area following 1 hour of CTS at 1 or 2 Hz (Fig. 1e); and nuclear area was significantly decreased following 1 hour CTS at 5 Hz (*p* = 0.003; Fig. 1f, Supplementary Fig. 1f), recovering after 24 hours. Under all CTS conditions, ratios of nuclear to cytoplasmic area were significantly decreased immediately following strain (*p* < 0.05; Supplementary Fig. 1g). Thus CTS was found to decouple the coordinated behaviour of cell and nuclear spreading observed at equilibrium on stiffness-defined substrates, either through failure of the nucleus to match CTS-induced cellular spreading (CTS at 1 and 2 Hz), or through nuclear contraction while cell spreading remained constant (at 5 Hz; Fig. 1g). Dynamic loading was thus accompanied by a disruption of the mechanisms linking the cyto- and nucleoskeletons.

### CTS-induced nuclear contraction requires stretch-activated ion channels

Stretch activated ion-channels can enable rapid response to mechanical stimulation^30^. To investigate the role of ion channels in CTS-induced nuclear contraction, we combined high-intensity strain with a panel of ion channel inhibitors: GdCl_3_, a broad-spectrum inhibitor of stretch-activated ion channels^31^; RN9893, an inhibitor of transient receptor potential cation channel subfamily V member 4 (TRPV4)^32^; amiloride, an inhibitor of acid sensing ion channels (ASICs)^31^, and GsMTx4, an inhibitor of piezo channels^33^ (Fig. 2a, Supplementary Fig. 2a). GdCl_3_ inhibited nuclear contraction following strain, as did the TRPV4-specific drug RN9893 and, to a lesser extent, amiloride. GsMTx4 did not prevent nuclear contraction, although earlier work has shown it to be effective in inhibiting chromatin condensation under milder loader regimes (3% uniaxial strain at 1 Hz)^33^. This suggested that activation of different ion channels may be specific to the loading regime. Following all treatments, nuclear area was recovered to control levels 24 hours after straining (Fig. 2b).

**Figure 2.**
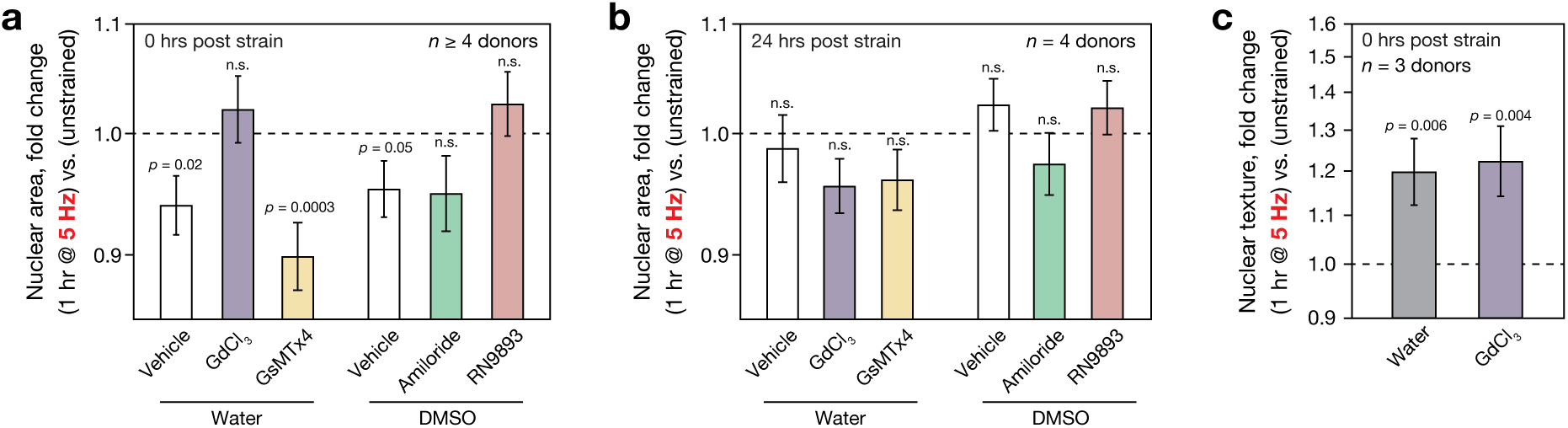
Ion channels are necessary for CTS-induced nuclear contraction, but not for chromatin condensation. (**a**) Changes to nuclear areas in primary hMSCs immediately following CTS (1 hour, 2.6 – 6.2% strain at 5 Hz; *n* ≥ 4 donors, see Supplementary Fig. 2a for an examination of variation between donors), normalized to unstrained controls. Ion channel activity was inhibited using: 10 μM GdCl_3_, a broad spectrum inhibitor of stretch-activated ion channels^31^; 3 μM GsMTx4, a specific inhibitor of piezo channels^33^; 100 μM amiloride, an inhibitor of acid-sensing ion channels (ASICs)^31^; 10 μM RN9893, a specific inhibitor of transient receptor potential vanilloid type 4 (TRPV4) channels^32^. Effects were compared to water or DMSO vehicle controls. Strain-induced reduction in nuclear area was prevented by inhibition of stretch-activated ion channels (GdCl_3_ inhibitor), specifically TRPV4 (RN9893 inhibitor). (**b**) Nuclear areas of hMSCs recovered to pre-strained values 24 hours after CTS, with no significant residual effect from ion channel inhibition. (**c**) Changes to nuclear texture assessed in primary hMSCs immediately following CTS treatment (1 hour, 2.6 – 6.2%, 5 Hz; *n* = 3 donors, see Supplementary Fig. 2b for donor-to-donor variation), normalized to unstrained control. Nuclear texture has been used to parameterize the extent of chromatin condensation^33, 34^, demonstrated by condensation with bivalent ions (Supplementary Figs. 2c, d). Increases in nuclear texture parameter following strain were unaffected by GdCl_3_ treatment, suggesting CTS-induced nuclear texture changes occurred independently of stretch-activated ion channels.

High-intensity CTS significantly increased the texture parameter of nuclear DAPI staining (*p* = 0.006, Fig. 2c, Supplementary Fig. 2b), indicative of chromatin condensation^33, 34^ (comparable to the effect of divalent ions, Supplementary Figs. 2c, d). Treatment with GdCl_3_ at its IC_50_ of 10 μM^35^ did not prevent changes to DAPI-stain texture following CTS. This contrasted with earlier characterizations of milder loading regimes, where GdCl_3_ was found to block chromatin condensation, although the drug concentration was higher in this case^33^. Our finding indicated the robustness of the chromatin condensation response in cells subjected to high-intensity CTS, but also suggested that chromatin condensation and contraction of nuclear area could be caused by different mechanisms.

### Cellular responses to high-intensity CTS are driven at the protein level

Using an initial targeted gene approach, we determined that the transcriptional response of hMSCs to 1 hour low-intensity CTS (1 Hz, change in strain = 4.0%), included upregulation of genes associated with cytoskeletal remodelling: intermediate filament vimentin (*VIM*) and alpha-actin-2 (*ACTA2*); and with the ECM: collagen alpha-1 (I) chain (*COL1A1*) and collagen alpha-1 (II) chain (*COL2A1*) (Supplementary Fig. 3a-d). These results are consistent with observed changes to cell morphology, and earlier characterizations of cellular responses to strain^26, 36^ and substrate stiffness^1^, which were proposed to increase robustness to stress. However, we found changes to the transcriptome assessed by RNA-Seq immediately following 1 hour high-intensity CTS to suggest only a narrow degree of change (Gaussian width = 0.21; Fig. 3a). Furthermore, gene ontology (GO) term analysis^37, 38^ of the genes affected by CTS suggested a general suppression of transcription and metabolism (Fig. 3b). Downregulation of transcriptional activity is consistent with our observations of chromatin condensation and previous reports of histone-methylation mediated gene silencing in endothelial progenitor cells subjected to low frequency (0.1 Hz) strain cycling^39^. The distribution of changes to gene expression was narrowed 24 hours after CTS (Gaussian width = 0.14; Supplementary Fig. 3e).

**Figure 3.**
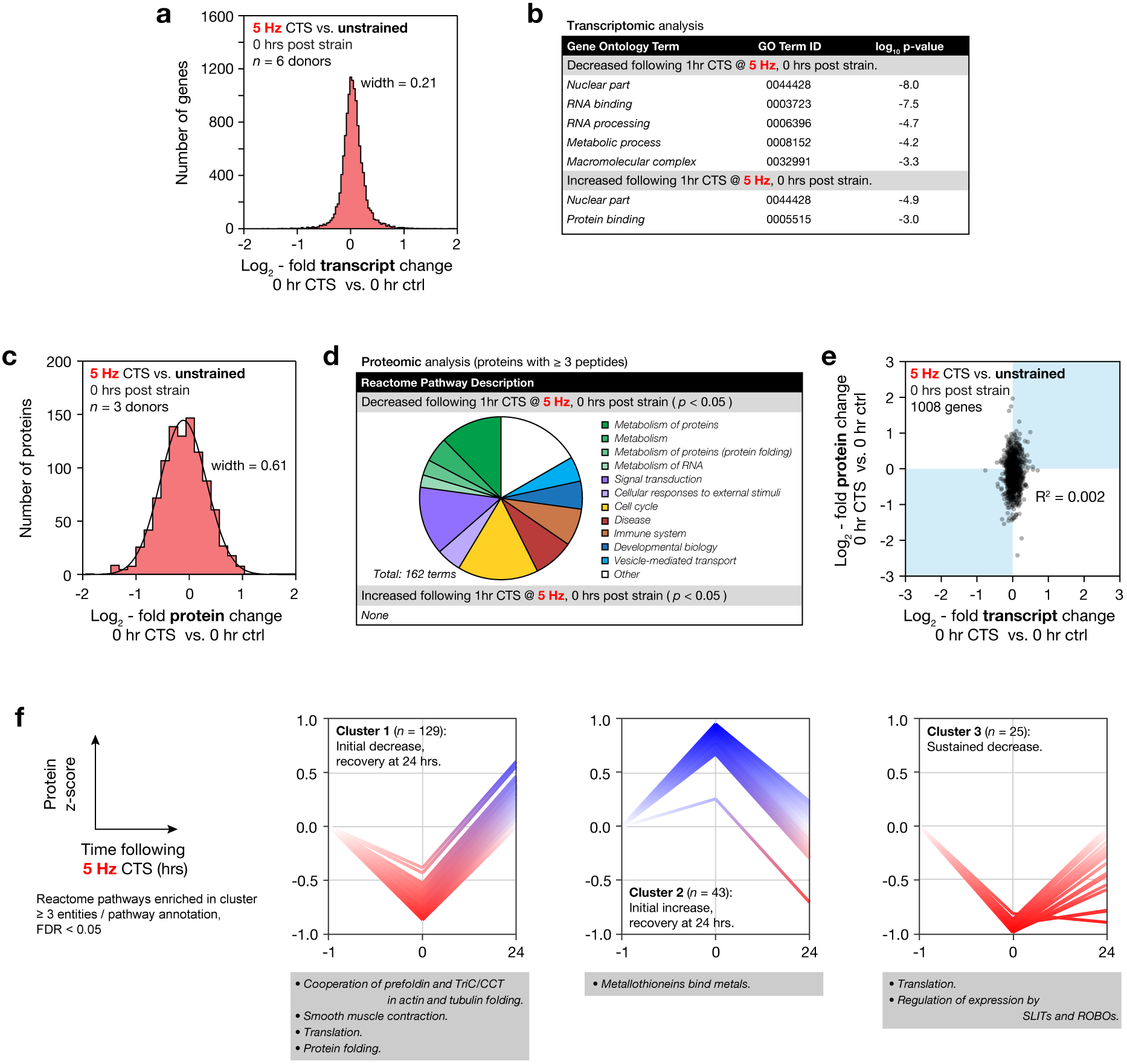
RNA processing and metabolism is down-regulated at the transcript-level following CTS, but there is a broad and uncorrelated protein-level response. (**a**) Histogram of changes to mRNA in primary hMSCs immediately following CTS (1 hour at 5 Hz, 2.6 – 6.2 % strain; *n* = 6 donors; data displayed as log_2_-fold transcript change CTS versus unstrained control). The width of a Gaussian curve fitted to the distribution was 0.21. (**b**) Gene ontology (GO) term analysis of the transcripts significantly up- or down-regulated following CTS. Annotations are suggestive of nuclear remodeling (nuclear component terms are both up- and down-regulated), and of down-regulation of RNA processing and metabolism, consistent with earlier studies in epidermal keratinocytes^39^. (**c**) Histogram of proteomic changes quantified by mass spectrometry (MS) immediately following CTS (1 hour at 5 Hz, 2.6 – 6.2 % strain; *n* = 3 donors; data displayed as log_2_-fold change following CTS, versus unstrained controls, and shows proteins quantified by three-or-more peptides; see Supplementary Fig. 3f for volcano plot). A Gaussian curve fit to the distribution had a width of 0.61, and was shifted to the left of zero (peak at -0.11 ± 0.01). (**d**) Analysis of Reactome pathways significantly affected at the protein level following CTS (*p* < 0.05, including proteins quantified by three-or-more peptides). Proteins associated with cellular metabolic processes were decreased following strain, including protein and RNA associated metabolism. Protein folding, signal transduction and external response pathways were also affected. (**e**) Correlation plot between proteome and transcriptome immediately following CTS (1008 genes quantified by RNA-seq and in proteomics by three-or-more peptides; R-squared = 0.002). The distribution of changes to protein levels was broader than – and uncorrelated with – transcript changes, suggesting regulation at the protein level. Transcript and protein levels were partially recovered 24 hours after CTS (Supplementary Figs. 3e, g-i). (**f**) K-means clustering was used to group quantified proteins based on their response to CTS after 0 and 24 hours. Analysis showed four possible clusters to be most appropriate for this dataset (Supplementary Fig. 3j). Three clusters are shown annotated with significant Reactome enrichments; ≥ 3 proteins per annotation; false discovery rate (FDR) < 0.05. RNA-Seq data can be found in Supplementary Table 1; MS proteomics data can be found in Supplementary Table 2.

In contrast to the analysis of transcript levels, analysis of the intracellular proteome quantified by mass spectrometry (MS), showed greater changes compared to unstrained controls (Fig. 3c, Supplementary Fig. 3f). A Gaussian fit to the distribution of protein fold changes had a width of 0.61 and was displaced to the left, suggesting that for an increased number of proteins, the rate of turnover was greater than the rate of translation. Analysis of the Reactome pathways^40-42^ significantly affected (false discovery rate (FDR)-corrected *p* < 0.05) by CTS showed that ontologies relating to metabolism of both protein and RNA, signal transduction and response to external stimuli were downregulated (Fig. 3d). Changes to transcript and proteome following CTS were not correlated (R-squared = 0.002; Fig. 3e), indicating a post-transcriptional regulation of protein levels. The proteome returned towards the control state after 24 hours (Gaussian width = 0.25; Supplementary Figs. 3g-i).

The time-resolved proteomic response to high-intensity CTS was further classified by K-means clustering (Fig. 3f, Supplementary Fig. 3j). Clusters of protein levels were identified with: (i) an immediate but unsustained decrease (cluster 1), enriched for Reactome annotations associated with translation, protein folding and mechanisms of actin and tubulin folding; (ii) an initial but unsustained increase (cluster 2), enriched for an annotation of metallothionein binding (associated with the management of oxidative stress^43^); and (iii) an immediate and sustained suppression (cluster 3), with enrichment of annotations for translation and regulation of the Slit/Robo signaling pathway (associated with cell polarity and cytoskeletal dynamics^44^). Taken as a whole, this analysis shows a complex, time-resolved and structured protein-level response to cellular stress management.

### CTS causes changes to protein conformation and post-translational modification

As changes in protein conformation are important to mechanotransduction^14, 15^, MS was performed following protein labeling with monobromobimane (mBBr), which by selectively labeling solvent-exposed Cys acts as an indicator of protein folding (Fig. 4a). MS was used to both identify mBBr-labeled proteins and quantify differential labeling in hMSCs following high-intensity CTS, relative to unstrained controls (Fig. 4b). The histogram of log_2_-fold changes in mBBr labeling showed a broad distribution of CTS-induced changes to mBBr reactivity, with labeling increased on average immediately following strain. The distribution was narrowed and centred about zero 24 hours after CTS, indicating a recovery of protein folding. Earlier applications of mBBr labeling have been used to identify force-dependent unfolding of domains in spectrin^45, 46^, cytoskeletal proteins^47^ and nuclear lamin-A/C (LMNA)^1^. Labeling of Cys522 in the Ig-folded domain of LMNA was previously used to report on the deformation of isolated nuclei subjected to shear stress in a rheometer^1^. We found the labeling of Cys522 to be increased 1.1-fold immediately following strain (FDR-corrected *p* < 0.001). We correlated changes to mBBr-labeled cysteine site occupancy and changes to total quantities of the parent proteins (Fig. 4c). This analysis suggested a systematic link between CTS-induced changes to protein conformation and stability (rates of translation vs. turnover), both on average (Figs. 3c, 4b) and in some cases on a protein-by-protein basis (Fig. 4c).

**Figure 4.**
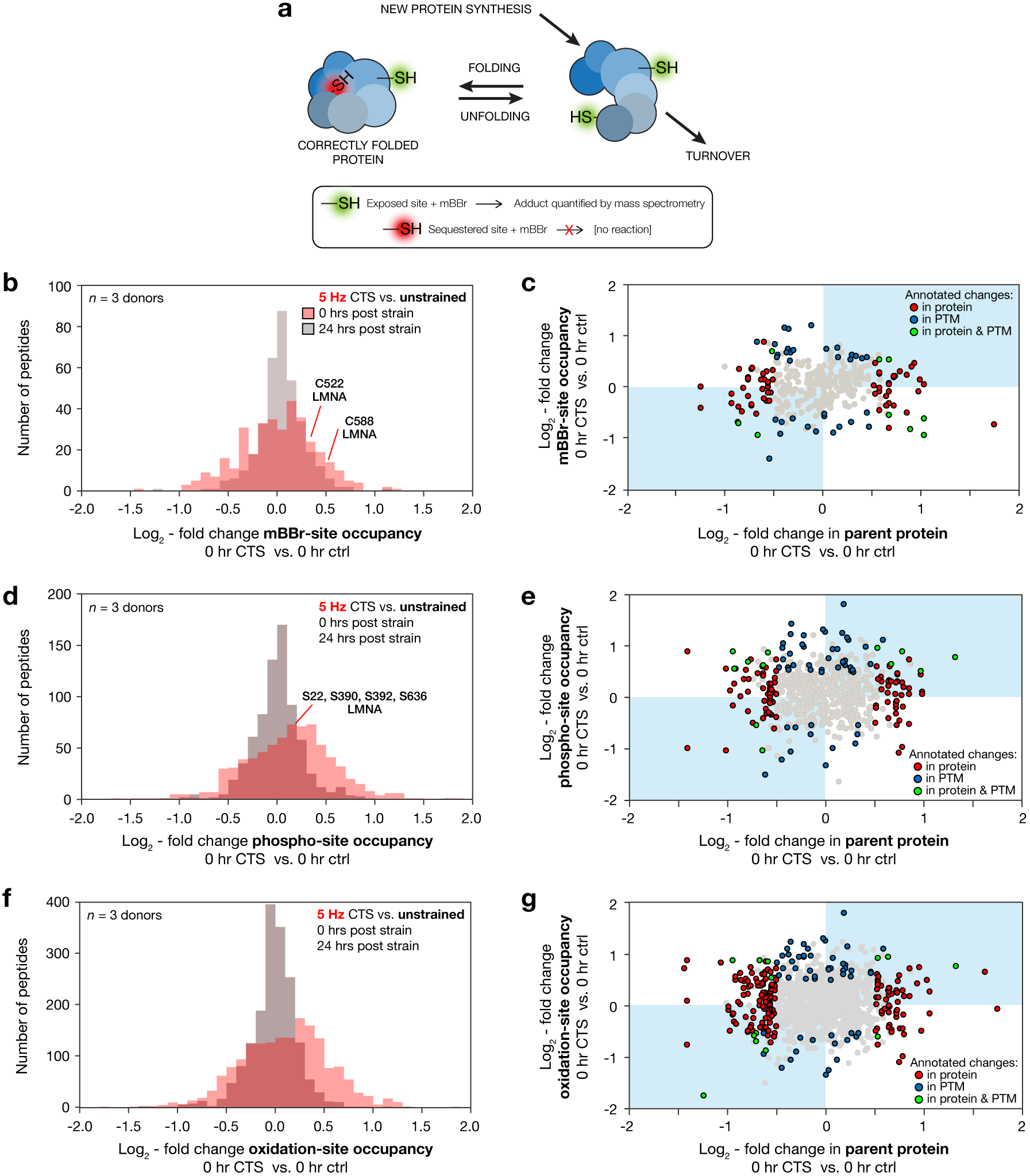
Changes in protein conformation and post translational modification (PTM) occur in concert with rapid regulation of protein levels in response to high-intensity CTS. (**a**) Schematic diagram of differential labeling of sulfhydryl groups by monobromobimane (mBBr)^1, 45^. Exposed cysteine residues are rapidly labeled, but those sequestered within folded proteins are unreactive; changes to the extent of labeling are indicative of altered protein conformation in response to a stress condition. Changes in protein quantity are determined by rates of synthesis versus turnover. (**b**) Histogram showing distribution of log_2_-fold changes to mBBr labeling site occupancy immediately and 24 hours after hMSCs were subjected to CTS (1 hour at 5 Hz, 2.6 – 6.2 % strain; *n* = 3 donors), measured relative to unstrained controls. The distribution immediately following CTS was broad and displaced to the right, indicating increased labeling; following 24 hours of recovery, the distribution was narrowed and centred around zero. Annotations indicate two sites within lamin-A,C (LMNA) with significantly increased modification (false discovery rate (FDR) corrected p-value < 0.05). Cys522 in LMNA was previously reported as a site of strain-sensitive mBBr labeling^1^. (**c**) Correlation between changes to mBBr labeling site occupancy and the quantity of the parent protein (i.e. source of the labeled peptide), immediately following CTS. In a number of cases, changes in mBBr modification, interpreted as a change in protein conformation or aggregation state, accompanied a change in the level of that protein. (**d**) Histogram showing changes to phospho-site occupancy immediately and 24 hours after CTS (1 hour at 5 Hz, 2.6 – 6.2 % strain; *n* = 3 donors), measured relative to unstrained controls. All phosphorylation sites shown have been curated previously in the PhosphoSitePlus database^70^. The distribution was shifted to the right immediately following CTS, indicating increased phosphorylation. (**e**) Correlation between phosphorylation and protein quantity. (**f**) Histogram showing changes to oxidation-site occupancy immediately and 24 hours after CTS (1 hour at 5 Hz, 2.6 – 6.2 % strain; *n* = 3 donors), measured relative to unstrained controls. (**g**) Correlation between oxidation and protein quantity; in 33% of cases, increased oxidation correlated with decreased protein levels (points in the top left quadrant). In figures (c), (e) and (g), data points are annotated as indicated in the legend if |log_2_-fold change| > 0.5 and the FDR corrected p-value < 0.05, otherwise they are shown in grey. Errors determined from linear modeling. MS proteomics data can be found in Supplementary Table 2.

Changes to endogenous PTMs were quantified by MS in the same experiment. A histogram of log_2_-fold changes to phospho-site occupancy following high-intensity CTS versus unstrained controls (Fig. 4d) showed increased phosphorylation. Phosphorylation of LMNA has been shown to be regulated in response to changes in substrate stiffness^1, 7^ and here we detected modest (~1.1-fold) but significant (FDR-corrected *p* < 0.001) increases in phosphorylation at S22, S390, S392 and S636. A correlation plot between changes of phosphorylation at individual residues and changes in quantities of the phosphorylated protein (Fig. 4e) showed that changed phosphorylation often accompanied modulation of protein quantity. A similar analysis of oxidised peptides showed that oxidation was increased immediately following CTS (Fig. 4f) and in a third of detected sites of oxidation, increased oxidation correlated with decreased protein levels (points in the top left quadrant of Fig. 4g). Previous work has established that cyclic stretching can increase levels of reactive oxygen species (ROS) in a range of cell types through activation of nicotinamide adenine dinucleotide phosphate (NADPH) oxidase systems and mitochondria. ROS have an important role in signal transduction, for example during vascularization, but can contribute to oxidative damage to lipids, proteins and DNA^48^. This potential for oxidative damage is perhaps consistent with the upregulation of protective metallothioneins (Fig. 3f). Profiles of both phosphorylation and oxidation became similar to controls 24 hours after CTS (Figs. 4d, f).

### CTS disrupts the linker of nucleo- and cytoskeleton (LINC) complex

Having established systematic responses to CTS, we sought to identify specific cases where protein regulation modulated mechano-transmission to the nucleus. There is a continuous system of structural proteins that connect cell interactions with the ECM at FAs, through the cytoskeleton and LINC complex to chromatin^13, 14, 20^. A central feature of this pathway is the LINC complex, which spans the nuclear envelope (NE) and includes nesprin proteins which bind to cytoskeletal components in the cytoplasm, but have Klarsicht, ANC-1, and Syne homology (KASH) domains extending into the nuclear lumen. The KASH domains bind to the Sad1 and UNC-84 (SUN) domains of SUN-domain containing proteins, which in turn bind to the nuclear lamina that lines the inner NE^49^. The nuclear lamina is composed of intermediate filament lamin proteins that confer structural integrity to the nucleus^2, 50, 51^ and also interface with chromatin and a range of regulatory and NE associated proteins^1, 52^. The complete system of protein linkages enables nuclear positioning^53^ and acts as a conduit for mechanical signals to regulate the genome^20, 52^.

LINC and NE protein levels were quantified by MS following 1 hour of high-intensity CTS, relative to unstrained controls (Fig. 5a). SUN-domain containing protein 2 (SUN2) was reduced to 52% of control levels (FDR-corrected *p* < 0.0001). In contrast, LMNA levels were not significantly altered. We also quantified proteins located specifically at the NE using immunofluorescence (IF) imaging (Fig. 5b, c, Supplementary Figs. 4a-e). IF confirmed that SUN2 levels were decreased at the NE following CTS (*p* = 0.03). Emerin (EMD), which has a role in the mechanical stimulation of the serum response factor (SRF) pathway^54^ was found to be significantly enriched at the NE (*p* < 0.0001), consistent with previous reports^39^. Additionally, we found that treatment with GdCl_3_ – shown to prevent contraction of nuclear area following CTS (Fig. 2a) – also prevented the loss of SUN2 (Fig. 5c, Supplementary Fig. 4f).

**Figure 5.**
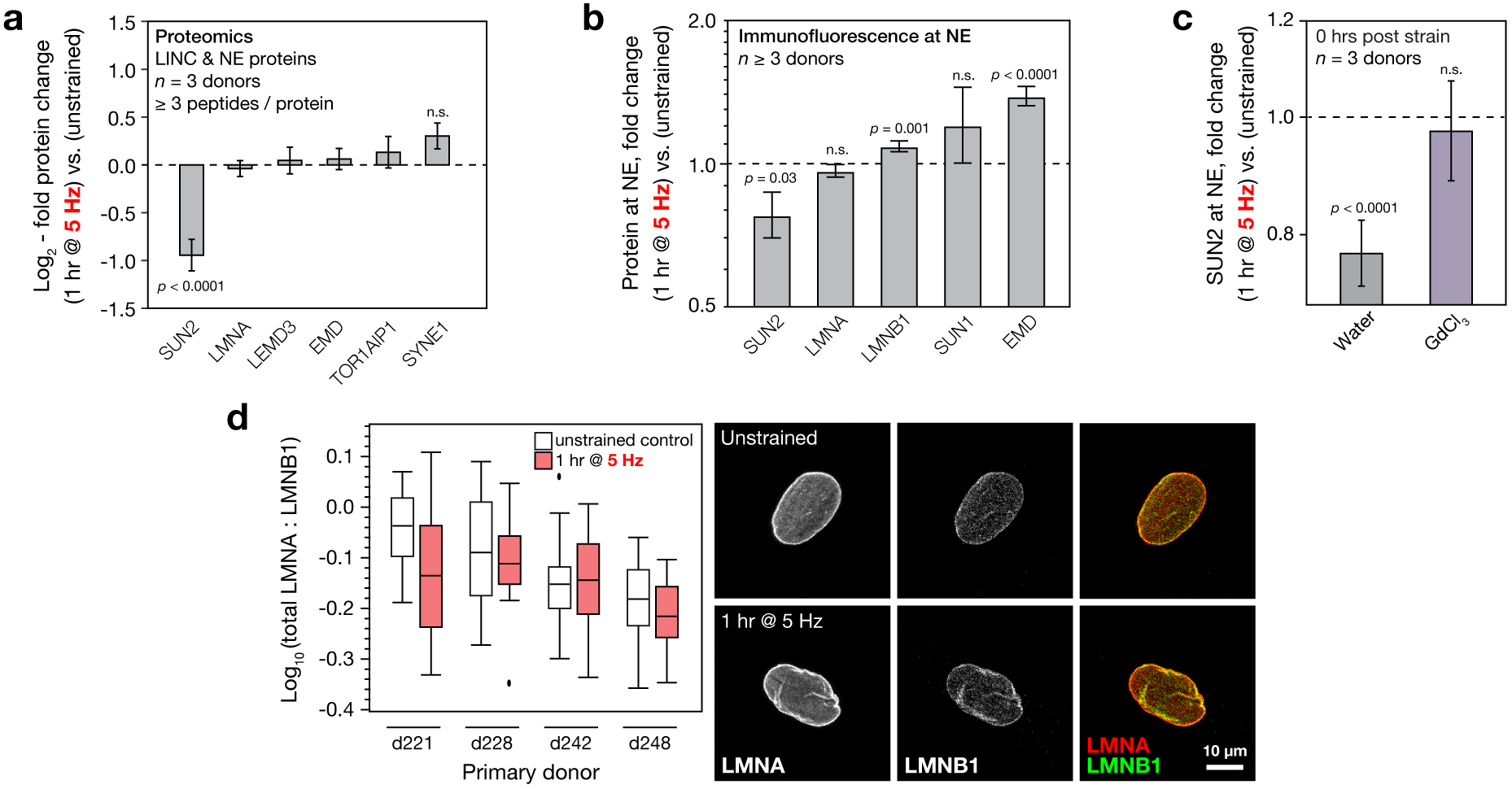
High intensity CTS affects the quantity of SUN domain-containing protein 2 (SUN2) and regulation of the linker of nucleo- and cyto-skeleton (LINC) complex. (**a**) Changes to LINC complex and nuclear envelope (NE) proteins in primary hMSCs, detected and quantified by MS immediately following CTS (1 hour at 5 Hz, 2.6 – 6.2 % strain; *n* = 3 donors), measured relative to unstrained controls. SUN2 was found to be significantly down-regulated (mean ± s.e.m. determined from linear modeling). (**b**) Immunofluorescence (IF) quantification of proteins localized at the NE immediately following CTS (1 hour at 5 Hz, 2.6 – 6.2 % strain; *n* ≥ 3 donors; see Supplementary Figs. 4a-e), using linear modeling to account for donor variation. (**c**) Changes to SUN2 levels measured by IF at the NE in hMSCs immediately following CTS (1 hour, 2.6 – 6.2% strain at 5 Hz; *n* = 3 donors, see Supplementary Fig. 4f for variation between donors) with ion channel inhibition (10 μM GdCl_3_), normalized to unstrained controls. Functional ion channels were required for significant down-regulation of SUN2 at the NE following CTS (mean ± s.e.m. determined from linear modeling). (**d**) Ratio of total lamin-A (LMNA) to lamin-B1 (LMNB1) quantified by IF in hMSCs immediately following CTS (*n* = 4 donors). CTS treatment decreased total LMNA:B1 to 0.92 ± 0.03 (p = 0.007; determined from linear modeling). Box-whisker plot shows means, quartiles, data spread and outliers determined by the Tukey method. Images show representative nuclei.

The composition of the nuclear lamina, characterised by the ratio of LMNA to lamin-B1 (LMNA:B1), has been used previously as a readout of nuclear adaption to the mechanical properties of the cellular microenvironment, with increased LMNA:B1 indicative of nuclear stiffening^1, 50^. We quantified LMNA:B1 by IF after 1 hour high-intensity CTS (Fig. 5d), finding it to be significantly decreased by CTS, relative to controls (*p* = 0.007); LMNB1 at the NE was significantly increased (*p* = 0.001). In contrast to the response of hMSCs to stiffer substrates^1^, the composition of the lamina was not altered to suggest stiffening following CTS.

### SUN2 is upstream of chromatin and cytoskeletal regulation

As decreased SUN2 at the NE was identified in response to CTS, we sought to further investigate its role. We quantified the effects of siRNA knockdown (KD) of SUN2 on primary hMSCs by MS proteomics, comparing two siRNAs to increase confidence of identifying on-target effects (Fig. 6a). The siRNAs showed KD of SUN2 to 35% of scrambled controls (Supplementary Figs. 5a, b). A Reactome pathway analysis of both KDs (Fig. 6b) identified significant perturbations to ‘*polycomb repressive complex 2 (PRC2) methylates histones and DNA*’, ‘*Protein lysine methyltransferases (PKMTs) methylate histone lysines*’ and ‘*Rho GTPases activate IQGAPs*’ (all FDR-corrected *p*-values < 0.05). These annotations suggested that SUN2 could be upstream of both aspects of chromatin regulation, consistent with our earlier observations, and nucleus-to-cytoskeleton (‘inside out’) signalling. A scatter plot of protein fold changes following SUN2 KD versus CTS showed correlation in cytoskeletal proteins such as actin and microtubule-associated protein (Fig. 6c), again suggesting that SUN2 regulation following CTS could be upstream of aspects of cytoskeletal remodelling.

**Figure 6.**
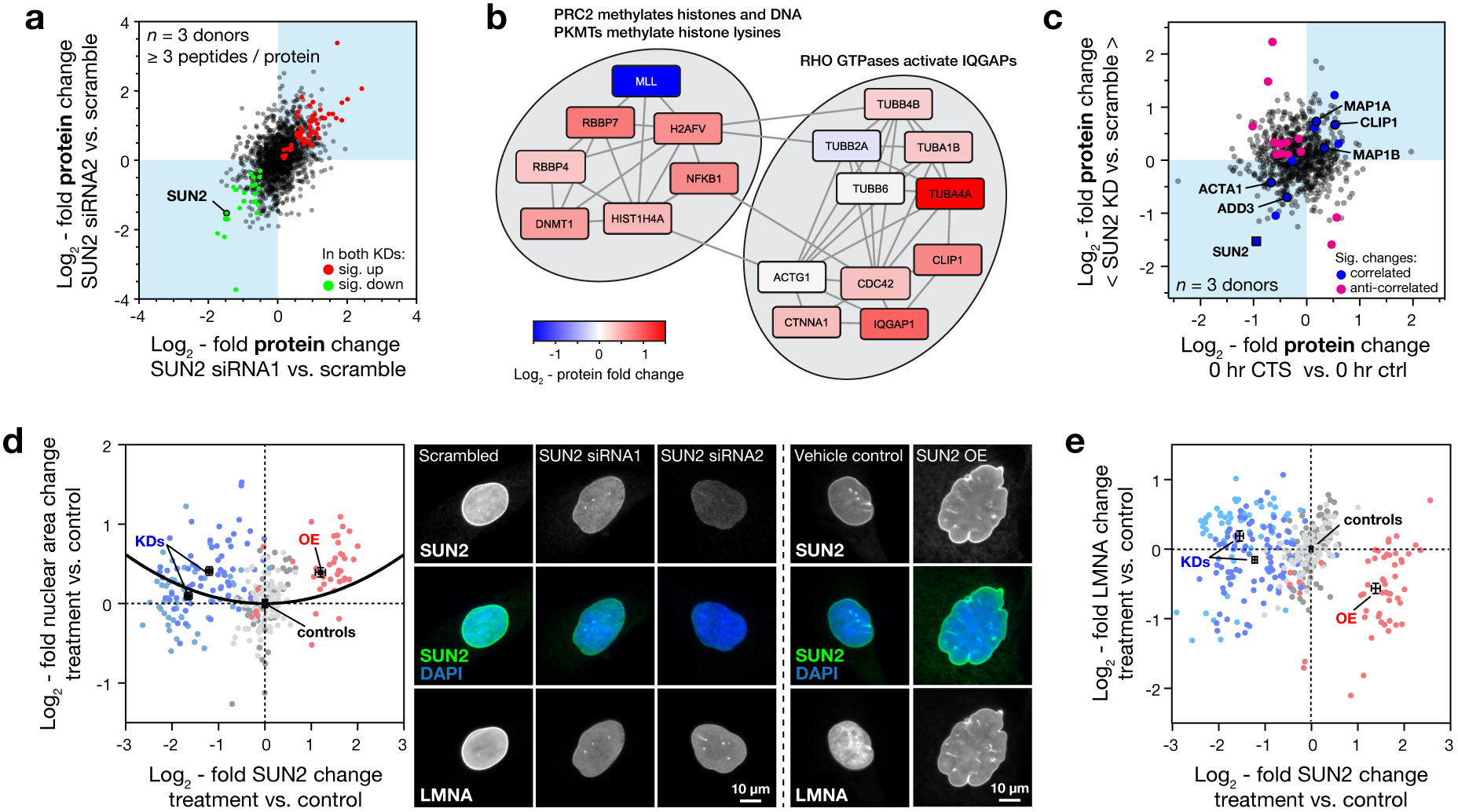
Knockdown (KD) of SUN2 affects the cellular proteome, cell and nuclear morphology. (**a**) MS characterization of the effects of SUN2 KD on protein levels in primary hMSCs. Plot shows correlation between log_2_-fold changes to the proteome following SUN2 KD with two separate siRNAs, siRNA1 and siRNA2, each measured relative to scrambled controls (quantification from at least three peptides-per-protein; *n* = 3 donors; see Supplementary Figs. 5a, b for histograms of fold changes). Data points annotated as indicated in the legend where significant changes occurred (false discovery rate, FDR, corrected p-value < 0.05), otherwise they are shown in grey. SUN2 was decreased by both siRNAs (to 35% of scrambled controls in both cases). (**b**) Reactome pathways significantly affected by both SUN2 KDs, relative to scrambled controls, shown with fold changes of constitutive proteins. (**c**) Correlation between changes to protein levels following SUN2 KD (mean of both siRNA treatments, relative to scrambled controls) and immediately following CTS (1 hour, 2.6 – 6.2% strain at 5 Hz relative to unstrained controls; see Fig. 3). Correlated and anti-correlated data annotated as indicated in the legend where significant changes occurred (FDR corrected p-value < 0.05; n = 3 donors; quantification based on three-or-more peptides-per-protein), otherwise they are shown in grey. (**d**) Changes to nuclear area in an immortalised hMSC line with SUN2 KD or overexpression (OE), normalised to controls. Nuclear areas were increased when SUN2 levels were perturbed from control levels (points represent measurements of individual cells; annotated black points show means for each condition; quadratic fit shown for guidance). See Supplementary Figs. 5c-h for data summaries and statistics. Images show representative nuclei stained for SUN2, lamin-A/C (LMNA) and DAPI. (**e**) Changes to LMNA levels in immortalised hMSCs with SUN2 KD or overexpression (OE), normalised to controls (see Supplementary Figs. 5k, l). MS proteomics data can be found in Supplementary Table 3.

### SUN2 modulates transmission of CTS to the nucleus and DNA damage

To investigate the role of SUN2 on mechano-transmission to the nucleus, we examined changes to nuclear morphology in immortalised hMSCs (Y201 line) shown to maintain the multipotency and mechano-responsiveness of primary MSCs^55, 56^. These were cultured on plastic with siRNA KD or doxycycline-induced overexpression (OE) of SUN2 (Fig. 6d, Supplementary Figs. 5c-h). In contrast to the response to CTS, where loss of SUN2 accompanied nuclear contraction, both KD and OE of SUN2 caused nuclear areas to increase. This suggested that there was a LINC complex composition determined by the cellular microenvironment – in this case, culture on plastic – and that imposed changes to protein levels caused a mismatch between nuclear and cellular properties^1^. We also found evidence supporting a disruption of nucleus-to-cytoskeleton signalling as both SUN2 KD and OE caused significantly increased cell spreading (Supplementary Figs. 5i, j). Consistent with an interpretation of LMNA as a reporter of a functioning mechanical linkage between the cytoskeleton and the nucleus, SUN2 OE led to loss of LMNA at the NE (Fig. 6e, Supplementary Figs. 5k, l).

Finally, we sought to determine how perturbation of SUN2 could affect cellular responses to CTS. We found that SUN2 KD in primary hMSCs was sufficient to prevent changes to nuclear:cytoplasmic area ratio following 1 hour of high-intensity CTS (Figs. 7a, b, Supplementary Figs. 6a-c). An siRNA treatment was also sufficient to block changes to nuclear texture indicative of chromatin condensation (Fig. 7c, Supplementary Fig. 6d). Likewise, SUN2 OE in immortalised hMSCs blocked the changes to nuclear:cytoplasmic area ratio observed in controls cells following 1 hour of high-intensity CTS (with recovery after 24 hours, Figs. 7d-f). Mechanical strain has previously been shown to cause DNA damage, inducing apoptosis in vascular smooth muscle cells^57^, and causing accumulation of damage to DNA and chromatin in nuclei subjected to extreme deformation as cells migrate through constricted environments^58-60^. We were surprised, therefore, to find that CTS here resulted in a small but significant decrease in the intensity of γH2AX staining in primary (*p* = 0.03) and immortalised MSCs (*p* = 0.0002), suggestive of a protective effect (Figs. 7g, h, Supplementary Fig. 6e). We found OE of SUN2 to override the decoupling response to CTS in immortalised hMSCs, concomitant with a significant increase in γH2AX staining (*p* < 0.0001; Fig. 7i). These results indicated that appropriate levels of SUN2 were essential for the mediation of nuclear decoupling in response to dynamic loading and therefore to afford protection to DNA.

**Figure 7.**
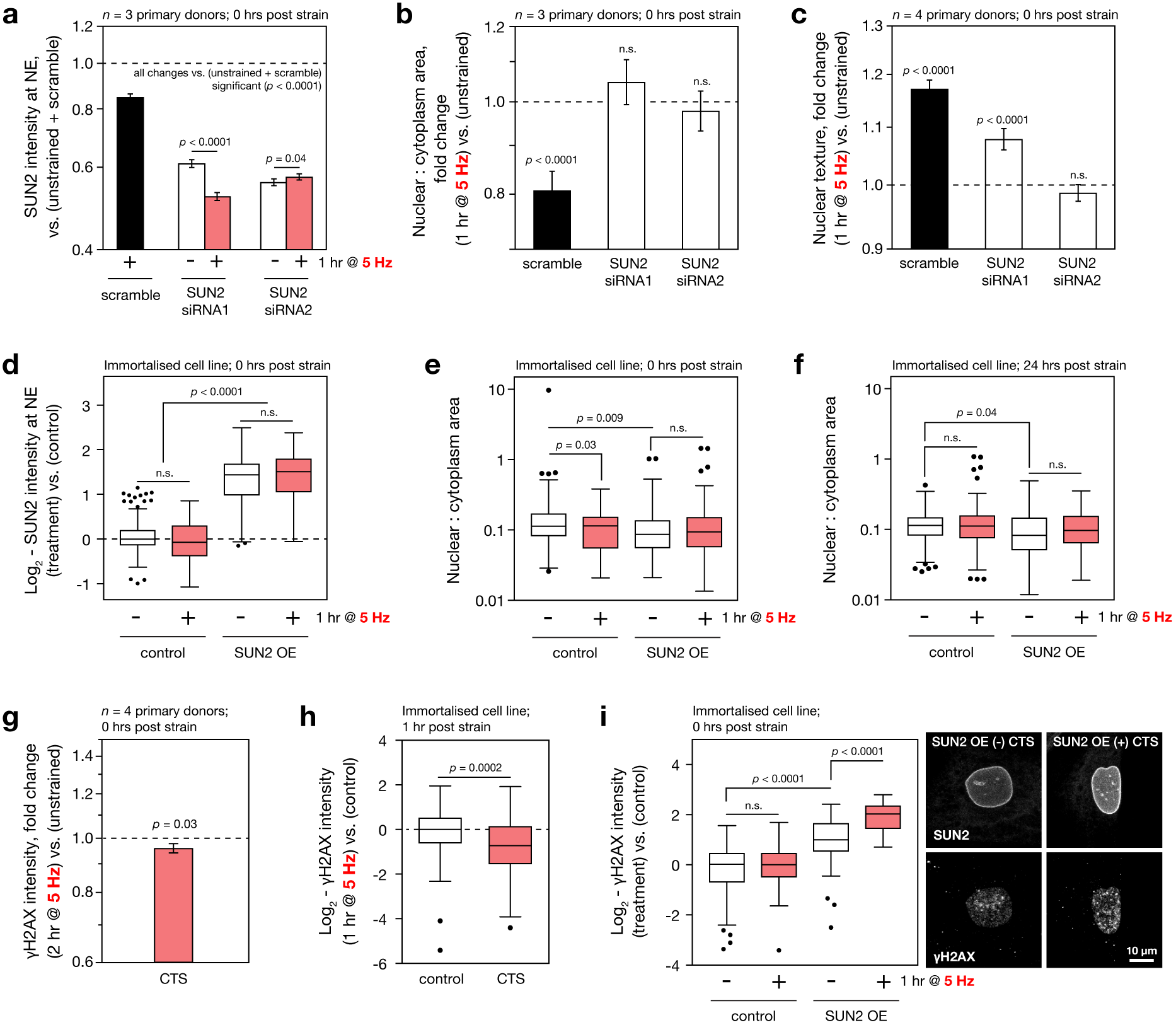
Both KD and OE of SUN2 block hMSC responses to CTS at the nucleus; γH2AX staining is increased in SUN2 OE hMSCs subjected to CTS. (**a**) SUN2 levels quantified by immunofluorescence (IF) at the nuclear envelope (NE) in primary hMSCs, following SUN2 KD (comparing effects of two siRNA sequences) and in combination with CTS (1 hour, 2.6 – 6.2% strain at 5 Hz). Both siRNAs were effective against SUN2 (see Fig. 6); SUN2 levels in the less potent KD (siRNA1) were further decreased by CTS, but the more efficient KD (siRNA2) showed little further change following CTS. (**b**) Ratios of nuclear to cytoplasmic areas in primary hMSCs following SUN2 KD and CTS. SUN2 KD with either siRNA sequence blocked the CTS-induced decrease to nuclear:cytoplasmic area characteristic of ‘nuclear decoupling’ (examined previously in Supplementary Fig. 1g). (**c**) Changes to chromatin texture in primary hMSCs following SUN2 KD and CTS. The more potent SUN2 KD (siRNA2) blocked strain-induced condensation of chromatin. Figure parts (a)-(c): *n* ≥ 3 donors; see Supplementary Figs. 6a-d for variation between donors; mean ± s.e.m. and *p*-values determined from linear modeling. (**d**) Quantification of changes to levels of SUN2 at the NE in immortalised hMSCs with inducible SUN2 OE, immediately following CTS (1 hour, 2.6 – 6.2% strain at 5 Hz). (**e**) Ratios of nuclear to cytoplasmic areas in immortalised hMSCs with SUN2 OE immediately following CTS. The nuclear:cytoplasmic area was significantly decreased in control cells following CTS, but the effect was blocked by overexpression of SUN2. (**f**) Differences in nuclear:cytoplasmic area ratios induced by CTS were lost 24 hours after straining. (**g**) The integrated intensity of γH2AX stained foci was significantly decreased in primary hMSCs immediately following CTS (2 hour, 2.6 – 6.2% strain at 5 Hz; *n* = 4 donors, see Supplementary Fig. S6e for donor-to-donor variation and representative images; mean ± s.e.m. and *p*-value determined from linear modeling). (**h**) γH2AX staining was also significantly decreased in immortalised hMSCs 1 hour after CTS (1 hour, 2.6 – 6.2% strain at 5 Hz; minimum of 98 cells per condition, *p*-value from t-test). (**i**) Quantification of γH2AX staining in immortalised hMSCs with inducible SUN2 OE, immediately following CTS (1 hour, – 6.2% strain at 5 Hz); CTS significantly increased DNA damage in cells with SUN2 OE. Figure parts (d)-(f), (i): Minimum of 27 nuclei analysed per condition; *p*-values determined from one-way ANOVA tests followed by Dunnett’s multiple comparison tests. Box-whisker plots show means, quartiles, data spread and outliers determined by the Tukey method. Images show representative nuclei.

## DISCUSSION

We have demonstrated in primary cells from multiple donors that hMSCs have a rapid, structured and reversible response to CTS regulated at the protein level. This response was dependent on both functional ion channels and appropriate levels of the LINC complex protein SUN2 (Fig. 8a). Furthermore, CTS was shown to cause changes within the LINC complex (Fig. 8b), in particular to the regulation of SUN2, enabling cells to decouple nuclear and cellular morphological behaviours and conferring protection to DNA. Robustness is increased through cyto- and nucleoskeletal remodeling in cells that have reached a ‘mechanical equilibrium’ state on increasingly stiff substrates^1, 61^. However, remodeling of the nuclear lamina seemed less important in the rapid response to high-intensity CTS. Although we were able to quantify modest (but significant) responses in LMNA conformation and phosphorylation, changes to the composition of the lamina were not indicative of nuclear stiffening. Mechano-transmission to the nucleus is an important mode of mechanical signaling, but if unregulated, has potential to apply stresses to chromatin. While a number of nuclear stress management mechanisms have been characterised, including chromatin condensation^33, 39^, chromatin detachment from the NE^62^, and altered nuclear mechanics^1, 7, 63^, a mechanism that isolates the nucleus from the cytoskeleton, as demonstrated here through regulation of SUN2, has potential to be both rapid and reversible. A role for SUN proteins in such mechanisms is further supported by analysis of protein turnover rates: SUN1 and 2 were reported to have the shortest half-lives of LINC complex proteins (Supplementary Fig. 7f)^64^.

**Figure 8.**
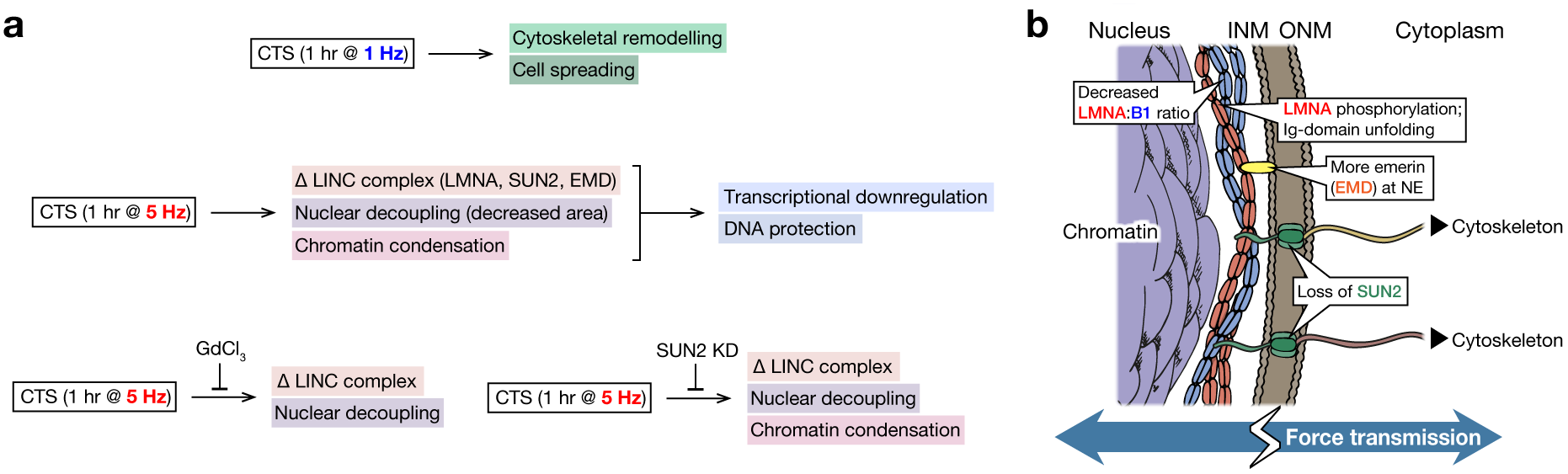
(**a**) Schematic summarizing the responses of mesenchymal stem cells (MSCs) to low and high intensity cyclic tensile strain (CTS), and how these pathways depend on ion channel activity and SUN domain-containing protein 2 (SUN2). (**b**) Cartoon summarizing strain-induced changes to proteins in the LINC complex and NE.

This study used large, unbiased ‘-omics’ datasets to identify and focus on regulation of the LINC complex. However, the techniques described here have potential to explore other aspects of the global cellular response to mechanical stress in greater detail. These include regulation of other structural proteins, such as the intermediate filaments^65^, molecular chaperones and the pathways that manage DNA and oxidative damage. The use of MSCs as a model system to study mechano-responsive processes has been widespread, but these cells are also being assessed for their potential for therapy in heart^19^ and muscle^18^ – tissues subject to sustained and high-frequency deformation^66^. Furthermore, this work may be particularly relevant to understanding how mechanical stress contributes to age-related pathology. Many aspects of the cellular stress response are abrogated in ageing^67^, but crucially, the NE may be particularly susceptible to misregulation^68, 69^.

### Methods

Methods and any associated references are available in the online version of the paper.

*Note: Supplementary Information is available in the online version of the paper.*

## Acknowledgements

HTJG and JS were funded by a Biotechnology and Biological Sciences Research Council (BBSRC) David Phillips Fellowship (BB/L024551/1). VM was partially supported by a studentship from the Sir Richard Stapley Educational Trust. OD was supported by a Wellcome Trust Institutional Strategic Support Fund (097820/Z/11/B). Proteomics was carried out at the Wellcome Trust Centre for Cell-Matrix Research (WTCCMR; 203128/Z/16/Z) Biological Mass Spectrometry Core Research Facility; RNA-Seq was performed by the Genomic Technologies Core Facility (GTCF). We thank Professor Paul Genever (University of York, UK) for use of the Y201 immortalised hMSC line; Professor Tim Hardingham (University of Manchester, UK) for useful discussions; Drs. Ronan O’Cualain and David Knight (WTCCMR) for advice on mass spectrometry analysis.

## Author Contributions

Investigation, HTJG, VM, OD, RP, MRJ and JS; Formal Analysis, HTJG, VM and JS; Writing – Original Draft, HTJG; Writing – Visualization, Review & Editing, HTJG, VM, OD, MRJ, RP, APG, SMR and JS; Project Administration and Funding Acquisition, JS.

## References

1. Swift, J. et al. Nuclear lamin-A scales with tissue stiffness and enhances matrix-directed differentiation. Science 341, 1240104 (2013).

2. Swift, J. & Discher, D.E. The nuclear lamina is mechano-responsive to ECM elasticity in mature tissue. J. Cell Sci. 127, 3005–3015 (2014).

3. Humphrey, J.D. Continuum biomechanics of soft biological tissues. Proc. R. Soc. A-Math. Phys. Eng. Sci. 459, 3–46 (2003).

4. Discher, D.E., Janmey, P. & Wang, Y.L. Tissue cells feel and respond to the stiffness of their substrate. Science 310, 1139–1143 (2005).

5. Levental, I., Georges, P.C. & Janmey, P.A. Soft biological materials and their impact on cell function. Soft Matter 3, 299–306 (2007).

6. Pritchard, R.H., Huang, Y.Y.S. & Terentjev, E.M. Mechanics of biological networks: from the cell cytoskeleton to connective tissue. Soft Matter 10, 1864–1884 (2014).

7. Buxboim, A. et al. Matrix elasticity regulates lamin-A,C phosphorylation and turnover with feedback to actomyosin. Curr. Biol. 24, 1909–1917 (2014).

8. Pelham, R.J. & Wang, Y.L. Cell locomotion and focal adhesions are regulated by substrate flexibility. Proc. Natl. Acad. Sci. USA 94, 13661–13665 (1997).

9. Lo, C.M., Wang, H.B., Dembo, M. & Wang, Y.L. Cell movement is guided by the rigidity of the substrate. Biophys. J. 79, 144–152 (2000).

10. Raab, M. et al. Crawling from soft to stiff matrix polarizes the cytoskeleton and phosphoregulates myosin-II heavy chain. J. Cell Biol. 199, 669–683 (2012).

11. McBeath, R., Pirone, D.M., Nelson, C.M., Bhadriraju, K. & Chen, C.S. Cell shape, cytoskeletal tension, and RhoA regulate stem cell lineage commitment. Dev. Cell 6, 483–495 (2004).

12. Engler, A.J., Sen, S., Sweeney, H.L. & Discher, D.E. Matrix elasticity directs stem cell lineage specification. Cell 126, 677–689 (2006).

13. Aureille, J., Belaadi, N. & Guilluy, C. Mechanotransduction via the nuclear envelope: a distant reflection of the cell surface. Curr. Opin. Cell Biol. 44, 59–67 (2017).

14. Humphrey, J.D., Dufresne, E.R. & Schwartz, M.A. Mechanotransduction and extracellular matrix homeostasis. Nat. Rev. Mol. Cell Biol. 15, 802–812 (2014).

15. Wasik, A.A. & Schiller, H.B. Functional proteomics of cellular mechanosensing mechanisms. Semin. Cell Dev. Biol. 71, 118–128 (2017).

16. Guilak, F. Biomechanical factors in osteoarthritis. Best Pract. Res. Clin. Rheumatol. 25, 815–823 (2011).

17. Phillip, J.M., Aifuwa, I., Walston, J. & Wirtz, D. The mechanobiology of aging. Ann. Rev. Biomed. Bioeng. 17, 113–141 (2015).

18. Turner, N.J. & Badylak, S.F. Regeneration of skeletal muscle. Cell Tissue Res. 347, 759–774 (2012).

19. Golpanian, S., Wolf, A., Hatzistergos, K.E. & Hare, J.M. Rebuilding the damaged heart: mesenchymal stem cells, cell-based therapy, and engineered heart tissue. Physiol. Rev. 96, 1127–1168 (2016).

20. Tajik, A. et al. Transcription upregulation via force-induced direct stretching of chromatin. Nat. Mater. 15, 1287–1296 (2016).

21. Engler, A. et al. Substrate compliance versus ligand density in cell on gel responses. Biophys. J. 86, 617–628 (2004).

22. Livne, A., Bouchbinder, E. & Geiger, B. Cell reorientation under cyclic stretching. Nat. Commun. 5, 8 (2014).

23. Wang, J.H.C., Goldschmidt-Clermont, P., Wille, J. & Yin, F.C.P. Specificity of endothelial cell reorientation in response to cyclic mechanical stretching. J. Biomech. 34, 1563–1572 (2001).

24. Hayakawa, K., Sato, N. & Obinata, T. Dynamic reorientation of cultured cells and stress fibers under mechanical stress from periodic stretching. Exp. Cell Res. 268, 104–114 (2001).

25. Smith, P.G., Garcia, R. & Kogerman, L. Strain reorganizes focal adhesions and cytoskeleton in cultured airway smooth muscle cells. Exp. Cell Res. 232, 127–136 (1997).

26. Greiner, A.M., Chen, H., Spatz, J.P. & Kemkemer, R. Cyclic tensile strain controls cell shape and directs actin stress fiber formation and focal adhesion alignment in spreading cells. Plos One 8, 9 (2013).

27. Elosegui-Artola, A. et al. Mechanical regulation of a molecular clutch defines force transmission and transduction in response to matrix rigidity. Nat. Cell Biol. 18, 540-+ (2016).

28. Buxboim, A. et al. Coordinated increase of nuclear tension and lamin-A with matrix stiffness outcompetes lamin-B receptor that favors soft tissue phenotypes. Mol. Biol. Cell 28, 3333–3348 (2017).

29. Maniotis, A.J., Chen, C.S. & Ingber, D.E. Demonstration of mechanical connections between integrins cytoskeletal filaments, and nucleoplasm that stabilize nuclear structure. Proc. Natl. Acad. Sci. USA 94, 849–854 (1997).

30. Matthews, B.D. et al. Ultra-rapid activation of TRPV4 ion channels by mechanical forces applied to cell surface beta 1 integrins. Integr. Biol. 2, 435–442 (2010).

31. Martinac, B. The ion channels to cytoskeleton connection as potential mechanism of mechanosensitivity. BBA - Biomembranes 1838, 682–691 (2014).

32. Wei, Z.L. et al. Identification of orally-bioavailable antagonists of the TRPV4 ion-channel. Bioorganic Med. Chem. Lett. 25, 4011–4015 (2015).

33. Heo, S.J. et al. Biophysical regulation of chromatin architecture instills a mechanical memory in mesenchymal stem cells. Sci. Rep. 5, 14 (2015).

34. Irianto, J. et al. Osmotic challenge drives rapid and reversible chromatin condensation in chondrocytes. Biophys. J. 104, 759–769 (2013).

35. Romanosilva, M.A., Gomez, M.V. & Brammer, M.J. The use of gadolinium to investigate the relationship between Ca2+ influx and glutamate release in rat cerebrocortical synaptosomes. Neurosci. Lett. 178, 155–158 (1994).

36. Gupta, V. & Grande-Allen, K.J. Effects of static and cyclic loading in regulating extracellular matrix synthesis by cardiovascular cells. Cardiovasc. Res. 72, 375–383 (2006).

37. Eden, E., Navon, R., Steinfeld, I., Lipson, D. & Yakhini, Z. GOrilla: a tool for discovery and visualization of enriched GO terms in ranked gene lists. BMC Bioinformatics 10, 7 (2009).

38. Supek, F., Bosnjak, M., Skunca, N. & Smuc, T. REVIGO summarizes and visualizes long lists of gene ontology terms. Plos One 6, 9 (2011).

39. Le, H.Q. et al. Mechanical regulation of transcription controls Polycomb-mediated gene silencing during lineage commitment. Nat. Cell Biol. 18, 864-+ (2016).

40. Croft, D. et al. The Reactome pathway knowledgebase. Nucl. Acids Res. 42, D472–D477 (2014).

41. Fabregat, A. et al. The Reactome pathway knowledgebase. Nucl. Acids Res., gkx1132–gkx1132 (2017).

42. Mallikarjun, V., Richardson, S.M. & Swift, J. Bayesian elastic nets for quantification of proteins and modifications in heterogeneous samples. BioRxiv, 10.1101/295527 (2018).

43. Ruttkay-Nedecky, B. et al. The role of metallothionein in oxidative stress. Int. J. Mol. Sci. 14, 6044–6066 (2013).

44. Ypsilanti, A.R., Zagar, Y. & Chedotal, A. Moving away from the midline: new developments for Slit and Robo. Development 137, 1939–1952 (2010).

45. Johnson, C.P., Tang, H., Carag, C., Speicher, D.W. & Discher, D.E. Forced unfolding of proteins within cells. Science 317, 663–666 (2007).

46. Krieger, C. et al. Cysteine shotgun-mass spectrometry (CS-MS) reveals dynamic sequence of protein structure changes within mutant and stressed cells. Proc. Natl. Acad. Sci. USA 108, 8269–8274 (2011).

47. Engler, A.J. et al. Embryonic cardiomyocytes beat best on a matrix with heart-like elasticity: scar-like rigidity inhibits beating. J. Cell Sci. 121, 3794–3802 (2008).

48. Birukov, K.G. Cyclic stretch, reactive oxygen species, and vascular remodeling. Antioxid. Redox. Sign. 11, 1651–1667 (2009).

49. Lombardi, M.L. et al. The interaction between nesprins and SUN proteins at the nuclear envelope is critical for force transmission between the nucleus and cytoskeleton. J. Biol. Chem. 286, 26743–26753 (2011).

50. Harada, T. et al. Nuclear lamin stiffness is a barrier to 3D migration, but softness can limit survival. J. Cell Biol. 204, 669–682 (2014).

51. Lammerding, J. et al. Lamins A and C but not lamin B1 regulate nuclear mechanics. J. Biol. Chem. 281, 25768–25780 (2006).

52. Gruenbaum, Y. & Foisner, R. Lamins: nuclear intermediate filament proteins with fundamental functions in nuclear mechanics and genome regulation. Annu. Rev. Biochem. 84, 131–164 (2015).

53. Luxton, G.W.G. & Starr, D.A. KASHing up with the nucleus: novel functional roles of KASH proteins at the cytoplasmic surface of the nucleus. Curr. Opin. Cell Biol. 28, 69–75 (2014).

54. Ho, C.Y., Jaalouk, D.E., Vartiainen, M.K. & Lammerding, J. Lamin A/C and emerin regulate MKL1-SRF activity by modulating actin dynamics. Nature 497, 507–511 (2013).

55. James, S. et al. Multiparameter analysis of human bone marrow stromal cells identifies distinct immunomodulatory and differentiation-competent subtypes. Stem Cell Rep. 4, 1004–1015 (2015).

56. Galarza Torre, A. et al. An immortalised mesenchymal stem cell line maintains mechano-responsive behaviour and can be used as a reporter of substrate stiffness. BioRxiv, 10.1101/269225 (2018).

57. Mayr, M., Hu, Y.H., Hainaut, P. & Xu, Q.B. Mechanical stress-induced DNA damage and rac-p38MAPK signal pathways mediate p53-dependent apoptosis in vascular smooth muscle cells. Faseb J. 16, 1423–1425 (2002).

58. Raab, M. et al. ESCRT III repairs nuclear envelope ruptures during cell migration to limit DNA damage and cell death. Science 352, 359–362 (2016).

59. Denais, C.M. et al. Nuclear envelope rupture and repair during cancer cell migration. Science 352, 353–358 (2016).

60. Irianto, J. et al. DNA damage follows repair factor depletion and portends genome variation in cancer cells after pore migration. Curr. Biol. 27, 210–223 (2017).

61. Solon, J., Levental, I., Sengupta, K., Georges, P.C. & Janmey, P.A. Fibroblast adaptation and stiffness matching to soft elastic substrates. Biophys. J. 93, 4453–4461 (2007).

62. Kumar, A. et al. ATR mediates a checkpoint at the nuclear envelope in response to mechanical stress. Cell 158, 633–646 (2014).

63. Guilluy, C. et al. Isolated nuclei adapt to force and reveal a mechanotransduction pathway in the nucleus. Nat. Cell Biol. 16, 376–381 (2014).

64. Schwanhausser, B. et al. Global quantification of mammalian gene expression control. Nature 473, 337–342 (2011).

65. Toivola, D.M., Strnad, P., Habtezion, A. & Omary, M.B. Intermediate filaments take the heat as stress proteins. Trends Cell Biol. 20, 79–91 (2010).

66. McAuley, J.H., Rothwell, J.C. & Marsden, C.D. Frequency peaks of tremor, muscle vibration and electromyographic activity at 10 Hz, 20 Hz and 40 Hz during human finger muscle contraction may reflect rhythmicities of central neural firing. Exp. Brain Res. 114, 525–541 (1997).

67. Kourtis, N. & Tavernarakis, N. Cellular stress response pathways and ageing: intricate molecular relationships. EMBO J. 30, 2520–2531 (2011).

68. Shimi, T. et al. The role of nuclear lamin B1 in cell proliferation and senescence. Genes Dev. 25, 2579–2593 (2011).

69. Scaffidi, P. & Misteli, T. Lamin A-dependent nuclear defects in human aging. Science 312, 1059–1063 (2006).

70. Hornbeck, P.V. et al. PhosphoSitePlus, 2014: mutations, PTMs and recalibrations. Nucl. Acids Res. 43, D512–D520 (2015).

